# An accessible metagenomic strategy allows for better characterization of invertebrate bulk samples

**DOI:** 10.1101/2025.01.17.633507

**Authors:** Martijn Callens, Guillaume Le Berre, Laure Van den Bulcke, Marianne Lolivier, Sofie Derycke

## Abstract

DNA-based techniques are a popular approach for assessing biodiversity in ecological research, especially for organisms which are difficult to detect or identify morphologically. Metabarcoding, the most established method for determining species composition and relative abundance in bulk samples, can be more sensitive and time- and cost effective than traditional morphological approaches. However, one drawback of this method is PCR bias caused by between-species variation in the amplification efficiency of a marker gene. Metagenomics, bypassing PCR amplification, has been proposed as an alternative to overcome this bias. Several studies have already shown the promising potential of metagenomics, but they all indicate the unavailability of reference genomes for most species in any ecosystem as one of the primary bottlenecks preventing its wider implementation. In this study, we present a strategy that uses unassembled reads of low-coverage whole genome sequencing to construct a genomic reference database, thus circumventing high sequencing costs and intensive bioinformatic processing. We show that this approach is superior to metabarcoding for approximating relative biomass of macrobenthos species from bulk samples. Furthermore, these results can be obtained with a sequencing effort comparable to metabarcoding. The strategy presented here can thus accelerate the implementation of metagenomics in biodiversity assessments, as it should be relatively easy to adopt by laboratories familiar with metabarcoding and can be used as an accessible alternative.

## INTRODUCTION

DNA-based methods are increasingly applied for biodiversity assessment and ecological monitoring. Currently, the most established method is metabarcoding, which relies on amplifying, sequencing, and classifying homologous taxonomic marker genes (for metazoans typically – but not limited to – COI or 18S) from bulk and environmental samples (Compson et al., 2020; Ruppert et al., 2019; Taberlet et al., 2012; van der Loos & Nijland, 2021). Metabarcoding can have several advantages over traditional morphology-based identifications such as an increased time- and cost effectiveness (Van den Bulcke et al., 2024), a reduced reliance on expert taxonomic knowledge (Shea et al., 2023), and a greater capacity for identifying cryptic species and juvenile stages (Jackson et al., 2014; Semmouri et al., 2021). However, its reliance on the PCR amplification of a taxonomic marker gene also creates some important and well-known drawbacks such as PCR amplification bias, caused by variation in hybridization efficiencies between the primers and templates originating from different species (Wintzingerode et al., 1997), and stochastic variation in template amplification (Kebschull & Zador, 2015). Low hybridization efficiencies for a particular species can result in systematic low amplification leading to an underestimation of species relative abundance, or even no amplification leading to false negatives. Concurrently, the relative abundance of other species is overestimated. There are several workarounds such as lowering annealing temperatures, the use of highly degenerate primers, or the use of a multi-locus approach targeting additional markers. However, these workarounds can be cumbersome, more expensive, or result in a higher degree of nonspecific amplification (Loos & Nijland, 2021) and do not address the problem of amplification stochasticity.

Several researchers have tried to overcome PCR amplification bias by using PCR-free approaches. Mitochondrial metagenomics, which relies on shotgun sequencing of community DNA and the subsequent taxonomic classification of mitochondrial DNA (mtDNA) reads, is a frequently explored PCR-free approach because of the relatively high number of available mitochondrial reference genomes (Arribas et al., 2016; Bista et al., 2018; Crampton-Platt et al., 2016; Tang et al., 2014). Because mtDNA generally represents less than 1% of the total DNA fraction, enrichment through e.g. centrifugation, hybrid-capture probes, or selective digestion of linear DNA is desirable to reduce the required sequencing depth (Macher et al., 2018; Ramón-Laca et al., 2023; Sevigny et al., 2021). This enrichment step, however, complicates DNA library preparation. Wagemaker et al. (2021) explored another PCR-free approach using genotype by sequencing on multispecies plant root samples (msGBS). Although successful, both DNA library preparation and analyses for msGBS still required labor-intensive steps such as digestion and calibration.

Probably the most straightforward PCR-free approach is direct shotgun sequencing and subsequent classification of all metagenomic DNA reads (Piñol, 2021). Indeed, several studies have already successfully used this approach on a range of community types (Bell et al., 2021; Cowart et al., 2018; Garrido-Sanz et al., 2020; Schmidt et al., 2022; Serite et al., 2023a). However, these studies point towards the lack of assembled reference genomes as a major bottleneck for applying this technique more widely. Ongoing ambitious large-scale reference genome projects are working towards filling this gap (Lewin et al., 2018; Mazzoni et al., 2023), but it might still take many years before a comprehensive list of reference genomes is available for most ecosystems.

To address the reference genome bottleneck and accelerate the wider application of PCR-free shotgun metagenomics on metazoan bulk samples, we developed a strategy to generate whole genome reference databases using unassembled low-coverage genome sequencing reads (Figure 1). We tested our strategy on shotgun metagenomic data from North Sea macrobenthos communities that were morphologically identified prior to bulk DNA extraction and for which we also obtained biomass and metabarcoding data. This allowed us to compare our metagenomic strategy to metabarcoding regarding the ability to capture community composition. We also performed extensive subsampling of the metagenomic sequencing data to provide recommendations for the required sequencing depths for both the generation of reference data and shotgun metagenomic sequencing of bulk samples.

**FIGURE 1:**
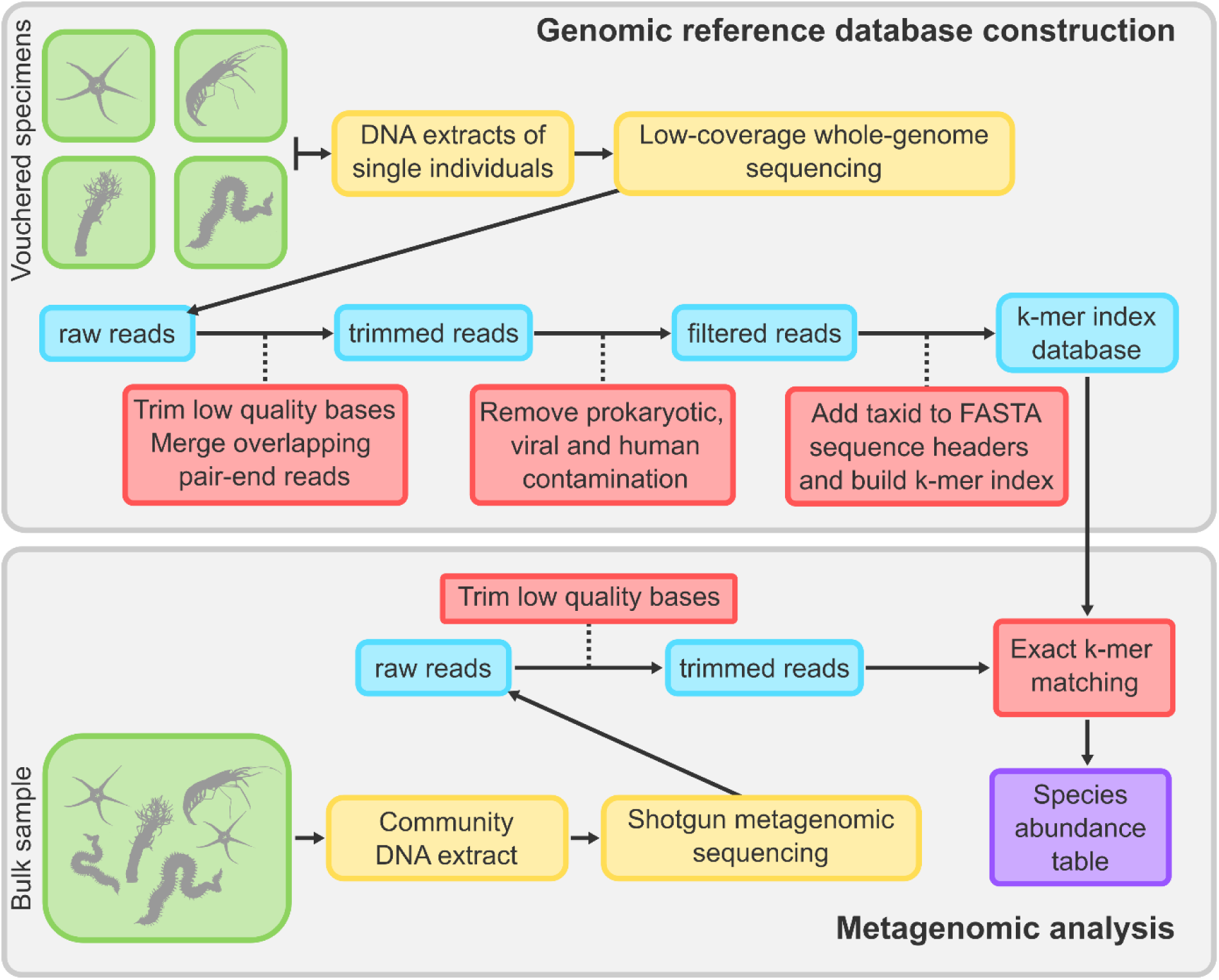
Schematic overview of the metagenomic strategy for characterization of invertebrate bulk samples. The top panel shows the workflow for assembly-free genomic reference database construction. The bottom panel shows the workflow for metagenomic analysis. Details for each step are described in the materials and methods section. Details and code for the bioinformatic processing steps (red squares) can be found in the supplementary materials.

## MATERIALS & METHODS

### Construction of a genomic reference database

#### Low-coverage genome sequencing of selected species

We selected 25 macrobenthos species for low-coverage genome sequencing to build our genomic reference database for metagenomic classification (Table 1). Macrobenthos species were selected based on three criteria: 1) species with a high relative abundance in one or several of our field samples based on morphological or metabarcoding data, 2) species for which we were unable to obtain a good reference COI sequence by sanger sequencing, and 3) species with a discrepancy in relative abundances based on morphological and metabarcoding data from our field samples. These criteria resulted in a set of species which allowed us to evaluate the performance of our PCR-free strategy (broad taxonomic distribution, differentiation in composition between field samples, and different types of correlations between morphological and metabarcoding data for the selected species). The commonly encountered European lancelet (*Branchiostoma lanceolatum*) was additionally included in our reference database using its published reference genome (genbank accession: GCF_035083965.1).

**TABLE 1.**
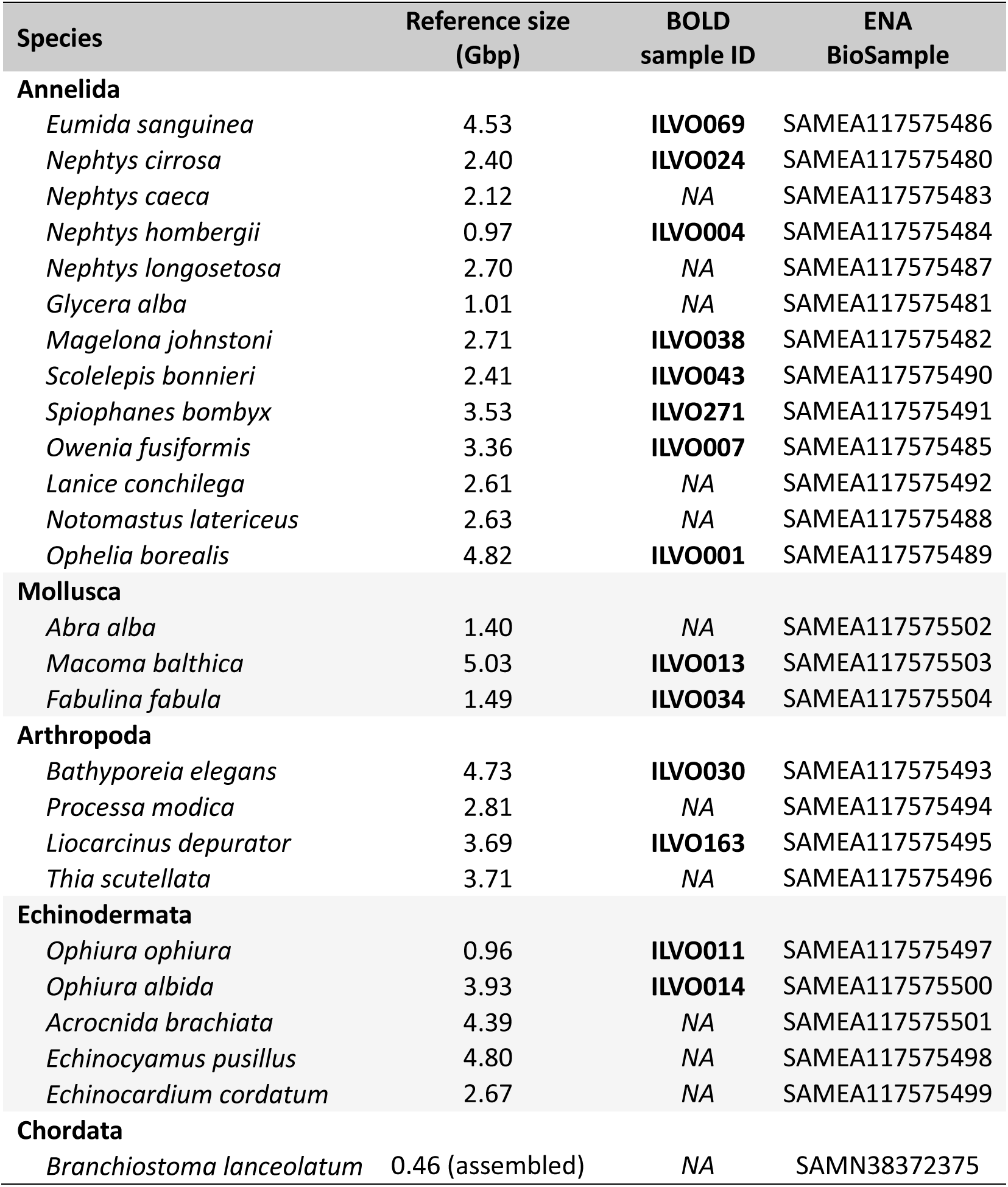
List of species included in this study. The reference size indicates the number of nucleotides for each species used to construct the custom kraken database. Vouchering information of species indicated with a BOLD sample ID is publicly available in BOLD (boldsystems.org). For species with no BOLD sample ID, no public data is available but specimen details have been recorded identical to those having a BOLD Sample ID and images are provided in the supplementary materials. Nucleotide data (raw reads and mitochondrial genome assemblies) are associated with the provided ENA BioSample, except for *B. lanceolatum* which is identified through an NCBI BioSample.

Genomic DNA of the selected macrobenthos species was obtained from our genomic reference material collection. This collection contains vouchered specimens of marine species mainly used for generating DNA barcodes (see Table 1 for BOLD Sample ID of the specimens used in this study and supplementary materials for specimen metadata). Genomic DNA was extracted using the Blood and Tissue kit (QIAGEN) and libraries for Illumina sequencing were prepared using the Illumina DNA prep kit (Illumina) and unique dual indexes for multiplexing according to the manufacturer’s instructions. For each sample, library insert sizes were checked on a bioanalyzer with a high-sensitivity DNA kit (Agilent), and DNA concentration was quantified with a Quantus fluorometer (Promega). Individual libraries were pooled, adjusting the volume of each library based on its DNA concentration. Pooled libraries were sequenced at the Oklahoma Medical Research Foundation NGS Core on an Illumina NovaSeq S4 flow cell (paired-end 2 x 150 bp), targeting 10 million paired-end reads per sample.

#### Bioinformatic processing of reference sequencing data

Sequencing quality of demultiplexed fastq files was checked using *FastQC* v0.12.1, and results from all fastq files were combined into a single report using *MultiQC* v1.13 (Ewels et al., 2016). Reads were quality trimmed with *Trimmomatic* v0.39 (Bolger et al., 2014) using the following parameters: ILLUMINACLIP:NexteraPE.fa:2:30:10, HEADCROP:3, TRAILING:3, SLIDINGWINDOW:4:30, MINLEN:36. Trimmed overlapping paired-end reads were merged using *FastqJoin* 1.3.1 (Aronesty, 2013), and combined with trimmed unpaired reads and non-overlapping read pairs into a single fastq file (considering each read from a non-overlapping read pair as a separate sequence). These fastq files were subsequently filtered for prokaryotic, viral and human contamination using *kraken2* (Wood et al., 2019) with the standard database (downloaded from https://benlangmead.github.io/aws-indexes/k2, compilation date 6/5/2023). Reads that remained unclassified (no matches against prokaryotic, viral or human genomes) were extracted to a new fasta file, and the NCBI taxonomic identifier of the species under consideration was added to the header of each sequence in this file.

After the sequencing data for all species was processed as described above, a custom kraken2 database was produced: all fasta files of the 26 species were added to a database library (*kraken2-build --add-to-library*), the NCBI taxonomy was downloaded (*kraken2-build --download-taxonomy*) and the database was build using default parameters (*kraken2-build --build*).

Additionally, we assembled the mitochondrial genomes of all sequenced species from quality-trimmed reads using GetOrganelle 1.7.7.0 (Jin et al., 2020). Assembled mitochondrial genomes were combined in a single fasta file, and this file was used to build a local BLAST database for the determination of a relative abundance threshold for removing potential false positives (see below).

### Metagenomic classification of community DNA

#### Field sampling and sample processing

Macrobenthos samples were collected by Van Veen grabs during two sampling campaigns on the Belgian part of the North Sea. Macrobenthos diversity in these samples was analyzed in earlier studies using morphological identification and metabarcoding (Derycke et al., 2021, 2023; Van den Bulcke et al., 2021, 2023). Details for sample collection and processing are described in Van Den Bulcke et al. (2021) for stations ZVL and 120; and in Van den Bulcke et al. (2024) for the 12 Thorntonbank (TB) stations. Briefly, at each station 0.1 m² Van Veen grabs were taken and sieved over a 1 mm sieve. The collected material was fixed in absolute ethanol and stored at −20°C until further processing. All macrobenthos specimens were morphologically identified and weighted prior to bulk DNA extraction. For all 14 samples analyzed in this study, metabarcoding and metagenomic analyses started from the same DNA extract. These DNA extracts can be directly linked to prior morphological identifications before extraction. The only exception is the ZVL sample, where the morphological identification was done on a separate replicate Van Veen sample simultaneously taken at the same location. The biomass of individuals morphologically identified as *Processa* sp. and *Glycera* sp. was respectively re-assigned to the species *Processa modica* and *Glycera alba*, as these were the most likely species using DNA barcoding and greatly improved agreement with both the metabarcoding- and metagenomic results.

Metagenomic DNA libraries for Illumina sequencing were simultaneously prepared with genomic DNA libraries for the genomic reference data using the same protocol, and metagenomic libraries were pooled with the genomic libraries on the same NovaSeq runs (2 x 150 bp, targeting 10 million paired-end reads per sample). A list of metagenomic samples along with their sequencing depth can be found in Supplementary Table S1.

#### Bioinformatic processing of metagenomic sequencing data

Raw read QC and trimming was done using the same methods as described above for reference sequencing data. The only difference was a lower required average quality within a 4 bp sliding window which was set to a Phred score of 15. After quality trimming, paired reads were classified using our custom kraken2 database with kraken 2.1.2, setting minimum-hit-groups to 3 (requires 3 hit groups to be found before declaring a sequence classified). After classifying all samples, the generated kraken reports were merged into a biom format using the kraken-biom tool (Dabdoub, 2016) and imported into R as a *phyloseq* object.

#### Setting a relative abundance threshold for removing potential false positives

Low abundance false positives are a known problem with exact k-mer matching (Breitwieser et al., 2018). This also seemed to affect our analysis, as taxonomic classification of metagenomic reads using exact k-mer matching generally resulted in the assignment of a small fraction of reads to the species that were not detected by morphology and metabarcoding. Therefore, we determined a relative abundance threshold for removing these potential false positives. This was done by assessing species presence in the metagenomic sample with the highest sequencing depth (TB54_19; 36.5 million reads) using an alignment-based method. This method for assessing species presence is much slower than k-mer matching and requires a high sequencing depth but was expected to be more accurate. We aligned metagenomic reads to assembled mitochondrial genomes using BLASTn (Camacho et al., 2009), and scored a species as present if at least 140 bp of at least one read uniquely aligned to the mitochondrial genome of this species with an identity greater than 97%. This analysis detected 10 species, and indicated that the fraction of reads assigned to potential false positives (i.e. species not detected by alignment to the mitochondrial genome) was generally less than 0.2%. Consequently, we considered a species as absent in a metagenomic sample when less than 0.2% of reads were assigned to that species.

### Similarity between morphology and DNA-based methods

#### Community composition – species detection and relative abundance

For biomass measurements and read counts obtained by metabarcoding, we only used data for species that were also included in our custom k-mer database (Table 1). For each sample, metabarcoding read counts less than 100 were removed to filter out potential false positives (Drake et al., 2022), which resulted in an overall better agreement of species detection between metabarcoding and morphological methods (Table 2). The relative biomass and the relative abundance of metabarcoding reads was calculated within each sample for each species using only this subset of data. For read counts obtained by metagenomics, we removed the read counts that were unclassified and read counts assigned to species that received less than 0.2% of the total amount of reads from each sample (to remove potential false positives, as explained above). This filtered dataset was then used to calculate the relative abundance of metagenomic read counts for each species per sample.

**TABLE 2.** Species detected in the same sample by a DNA-based methods compared to species detected by morphological identification and the resulting Jaccard similarity (J). Values between brackets indicate numbers before abundance filtering (removal of read counts < 100 reads for metabarcoding and removal of reads with a relative abundance < 0.2% for metagenomics). *For the ZVL sample morphological and DNA-based characterization were performed on separate replicate samples which might additionally explain discrepancies.

| Sample | Metabarcoding |  |  |  | Metagenomics |  |  |  |
| --- | --- | --- | --- | --- | --- | --- | --- | --- |
|  | Shared | Missing | Additional | J | Shared | Missing | Additional | J |
| 120 | 6 (9) | 6 (3) | 1 (6) | 0.46 | 12 | 0 | 2 (14) | 0.86 |
| ZVL* | 1 | 1 | 1 (3) | 0.33 | 1 (2) | 1 (0) | 1 (24) | 0.33 |
| TB11_19 | 2 | 0 | 3 (6) | 0.40 | 2 | 0 | 1 (24) | 0.67 |
| TB12_19 | 4 | 0 | 0 (3) | 1.00 | 3 (4) | 1 (0) | 2 (22) | 0.50 |
| TB12_21 | 5 | 0 | 0 (3) | 1.00 | 4 (5) | 1 (0) | 1 (21) | 0.67 |
| TB16_21 | 7 | 3 | 2 (7) | 0.58 | 8 (10) | 2 (0) | 2 (16) | 0.67 |
| TB47_21 | 4 | 0 | 0 (4) | 1.00 | 4 | 0 | 2 (22) | 0.67 |
| TB48_19 | 7 | 1 | 1 (4) | 0.78 | 7 (8) | 1 (0) | 2 (18) | 0.70 |
| TB48_21 | 4 | 0 | 2 (3) | 0.67 | 4 | 0 | 0 (22) | 1.00 |
| TB54_19 | 7 | 5 | 0 (3) | 0.58 | 10 (12) | 2 (0) | 1 (14) | 0.77 |
| TB54_21 | 8 (9) | 3 (2) | 2 (5) | 0.62 | 10 (11) | 1 (0) | 1 (15) | 0.83 |
| TB56_19 | 8 (9) | 4 (3) | 1 (3) | 0.62 | 8 (12) | 4 (0) | 2 (14) | 0.57 |
| TB59_19 | 7 | 3 | 2 (4) | 0.58 | 8 (10) | 2 (0) | 5 (16) | 0.53 |
| TB66_21 | 5 | 2 | 2 (7) | 0.56 | 7 | 0 | 5 (19) | 0.58 |

We used the Jaccard index to compare the species detected by metabarcoding or metagenomics to species detected by morphological identification within a sample. The Jaccard similarity (J) between a DNA based method and morphological identification was calculated for each sample as:

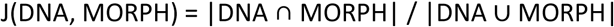

With |DNA ∩ MORPH| the number of species detected by both a DNA-based method and morphological identification, and |DNA ∪ MORPH| the total number of species detected with both methods. The statistical difference between Jaccard similarity values obtained by metabarcoding and metagenomics was tested using a non-parametric Wilcox test.

We used the Bray-Curtis similarity index to compare the community composition obtained by metabarcoding or metagenomics to the relative biomass-based community composition. For each sample, the Bray-Curtis similarity between relative read abundance and relative biomass was calculated for both metabarcoding and metagenomics. Given that we are dealing with relative abundances, the Bray-Curtis similarity can be calculated as: 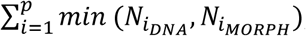; with *p* = the total number of observed species by both methods, *N*_*i*_*DNA*__ the relative read count of species *i* determined by a DNA-based methods (metabarcoding or metagenomics), and *N*_*i*_*MORPH*__ the relative biomass of species *i* determined by morphological identification. The statistical difference between Bray-Curtis similarity values obtained by metabarcoding and metagenomics was tested using a non-parametric Wilcox test. Differences between samples and methods were visualized by non-metric multidimensional scaling (NMDS) based on the Bray-Curtis distance.

#### Species-specific comparisons

PCR amplification bias often results in species-specific deviations between read abundance and biomass (caused by over- or under amplification). We investigated if metagenomics was able to alleviate this issue, especially for species with indications that a strong PCR bias exists. The assessment of the performance of both DNA-based methods was done by comparing our observed data with a simple linear regression model that assumes relative read counts have a one-to-one relation with relative biomass (y = x). Although other factors besides the precision of the method can cause deviations from this model (e.g. non-linear relationship between biomass and DNA content of an organism), a better fit to this model indicates a higher predictive value for relative biomass. We only included species in our analysis with biomass measurements in at least three samples (16 species in total). Goodness-of-fit to this model was determined by the coefficient of determination (R^2^). Given that our regression model was not derived from a model fitting procedure, R^2^ could take on negative values (which are reported as R^2^< 0). These negative values indicate that the overall mean of relative read abundance is a better predictor than the actual observations, which can be interpreted as the absence of predictive value for relative biomass.

### Estimations of required sequencing depth

#### Reference database

We estimated the minimal amount of genome sequencing data required per species to construct a custom kraken2 database with high read classification performance. This estimate was obtained by classifying metagenomic reads using kraken2 databases constructed from a range of sequencing depths. The number of classified reads with each database was then compared to the number of reads classified using a benchmark database constructed from an assembled reference genome. Because there were no species in our macrobenthos dataset for which we had both low-coverage genome sequencing reads and an assembled reference genome, we performed this analysis using two fish species (*Clupea harengus* and *Pleuronectes platessa*) for which both types of reference data were available from a previous project. Methods for low-coverage genome sequencing, custom kraken2 database construction and metagenomic classification were the same as those described earlier.

For each species, the benchmark kraken2 database was constructed using reference genomes obtained from NCBI RefSeq (*C. harengus*: GCF_900700415.2; *P. platessa*: GCF_947347685.1). We furthermore constructed kraken2 databases for both species using different random subsets of the reads from the low-coverage genome sequencing data (5 subsets containing 1, 2, 5, 10, 15 million reads, with extra subsets of 20 million reads for *C. harengus*). These subsets correspond to a sequencing depth and theoretical coverage of respectively the *C. harengus* and *P. platessa* genome of 0.3 Gbp (0.38X; 0.44X), 0.6Gbp (0.76X; 0.87X), 1.5Gbp (1.91X; 2.18X), 3Gbp (3.82X; 4.36X), 4.5Gbp (5.72X; 6.55X), and 6Gbp (7.63X; *NA*). Three replicate subsets were taken for each sequencing depth using *seqkit sample* (Shen et al., 2016). This resulted in a total of 17 and 20 kraken2 databases for *P. platessa* and *C. harengus*, respectively (1 benchmark database constructed using an assembled reference genome + 1 database from all unassembled reads + 3 × 5 or 6 subsets of unassembled reads).

For each fish species, we performed shotgun metagenomic sequencing on a positive control sample (consisting of eDNA extracted from seawater originating from a bucket in which multiple individuals of the fish species were present). The positive control metagenomic datasets for *P. platessa* and *C. harengus* contained 45.7 million and 1.4 million reads, respectively. Metagenomic reads were classified with all kraken2 databases for a particular species, and the fraction of reads classified with each database relative to the benchmark database was calculated.

#### Metagenomic sequencing

To estimate the effect of metagenomic sequencing depth on assessing community composition, we compared community composition obtained with subsets of metagenomic reads of the macrobenthos samples to community compositions obtained with the full set of metagenomic reads. For each metagenomic sample, subsets of 10^3^, 10^4^, 10^5^, 10^6^ and 10^7^ reads were produced in triplicate using *seqkit sample*. Subset of reads were classified using the same methods as described above for metagenomic samples. Results on community composition obtained with these subsets were compared to those obtained with the full set of metagenomic reads to calculate their Jaccard similarity and Bray-Curtis similarity.

## RESULTS

### Similarity between morphological and DNA-based methods

#### Species detection

We did not find a difference between the overall performance of metabarcoding and metagenomics for species detection. Compared to morphological identifications, there was no significant difference in Jaccard similarities obtained with metabarcoding and metagenomics (Wilcoxon rank sum test W = 108.5, p = 0.644; Figure 2A). The number of species shared between the morphology and both DNA-based methods was similar for metabarcoding and metagenomics (Table 2). Exceptions were stations 120, TB54_19, TB54_21, and TB66_21, where the metagenomic method found two to six extra shared species. Furthermore, stations TB11_19, TB12_19, TB47_21, TB59_19, and TB66_21 found two or three additional species with metagenomics compared to metabarcoding, while the opposite was true for TB48_21.

**FIGURE 2.**
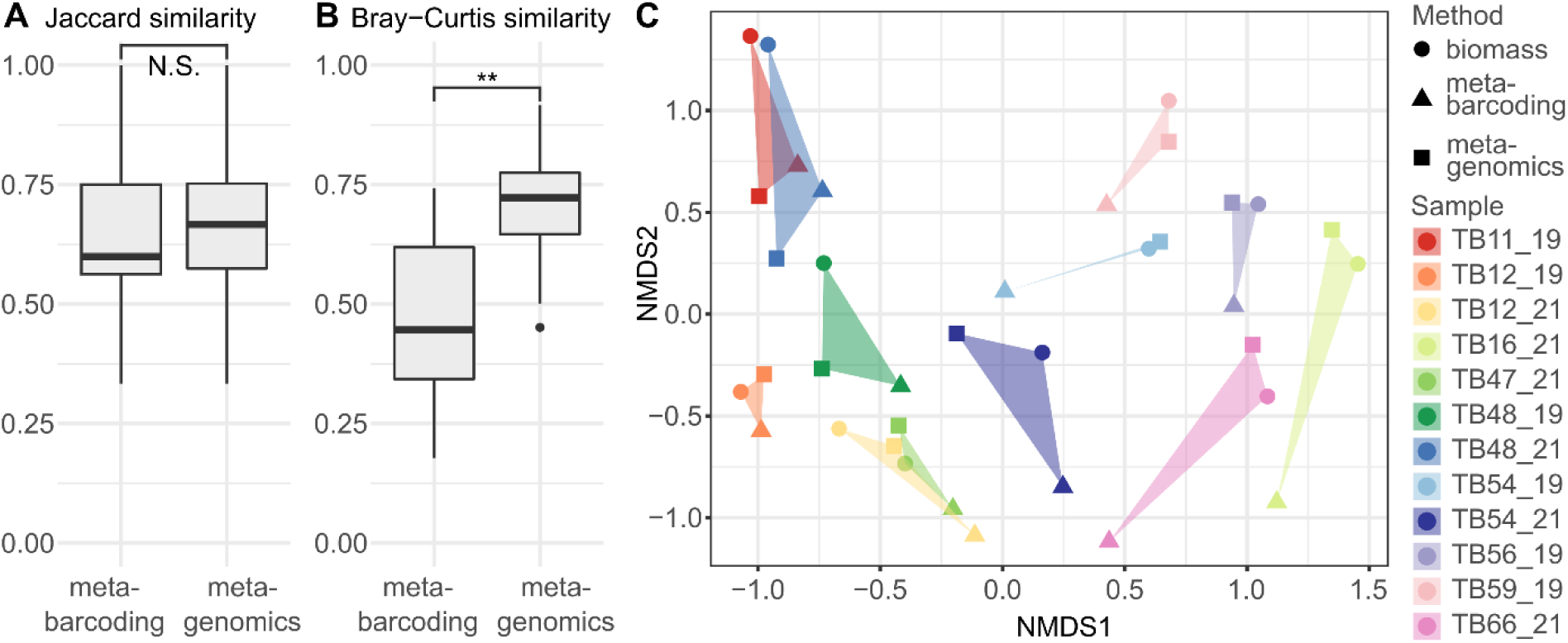
A) Jaccard similarity between DNA-based methods and morphological identification. B) Bray-Curtis similarity between DNA-based methods (relative read abundances) and morphological identification (relative biomass). ** significance level: p < 0.01. C) Nonmetric multidimensional scaling plot based on Bray-Curtis dissimilarity, samples are indicated by color, methods by point shape.

#### Relative abundance

We found that metagenomics performed significantly better than metabarcoding for assessing community composition. Community composition based on metagenomics had a higher Bray-Curtis similarity to community composition based on biomass for 9 out of the 12 samples. This resulted in a significantly higher overall Bray-Curtis similarity (Wilcoxon rank sum test W = 26, p = 0.007; Figure 2B and 2C). The only exceptions were three samples: TB11_19 with the same Bray-Curtis distance for both methods (both 0.73); and TB48_21, and TB48_19 with a lower Bray-Curtis distance for metagenomics than for metabarcoding (0.50 vs. 0.70 and 0.45 vs. 0.52, respectively). Interestingly, these three samples are characterized by a very high relative biomass of *Echinocardium cordatum* (TB11_19 = 99.1%, TB48_21 = 98.5% and TB48_19 = 52.7%).

#### Species-specific comparisons

We found that for 11 out of the 16 species analyzed, metagenomic relative read abundance had a better fit to relative biomass compared to metabarcoding. For three species, both metagenomic and metabarcoding relative read abundance had no predictive value for relative biomass (*Acrocnida brachiata, Eumida sanguinea,* and *Notomastus latericeus*). For two species, *E. cordatum* and *Owenia fusiformis*, metabarcoding relative read abundance had a better fit to relative biomass compared to metagenomics. For species where metagenomics provided a better fit, abundance-abundance plots indicate that metagenomics corrected for a negative PCR bias (under-amplification) in four species (Figure 3), and a positive PCR bias (over-amplification) in another four species (Figure 4). For the remaining three species where metagenomics provided a better fit, no clear pattern was discernible (Supplementary Figures S1 and S2). In addition, melting temperatures (T_m_) for the metabarcoding primers indicated a strong mismatch for most species which remained undetected by metabarcoding (Supplementary Table S2).

**FIGURE 3.**
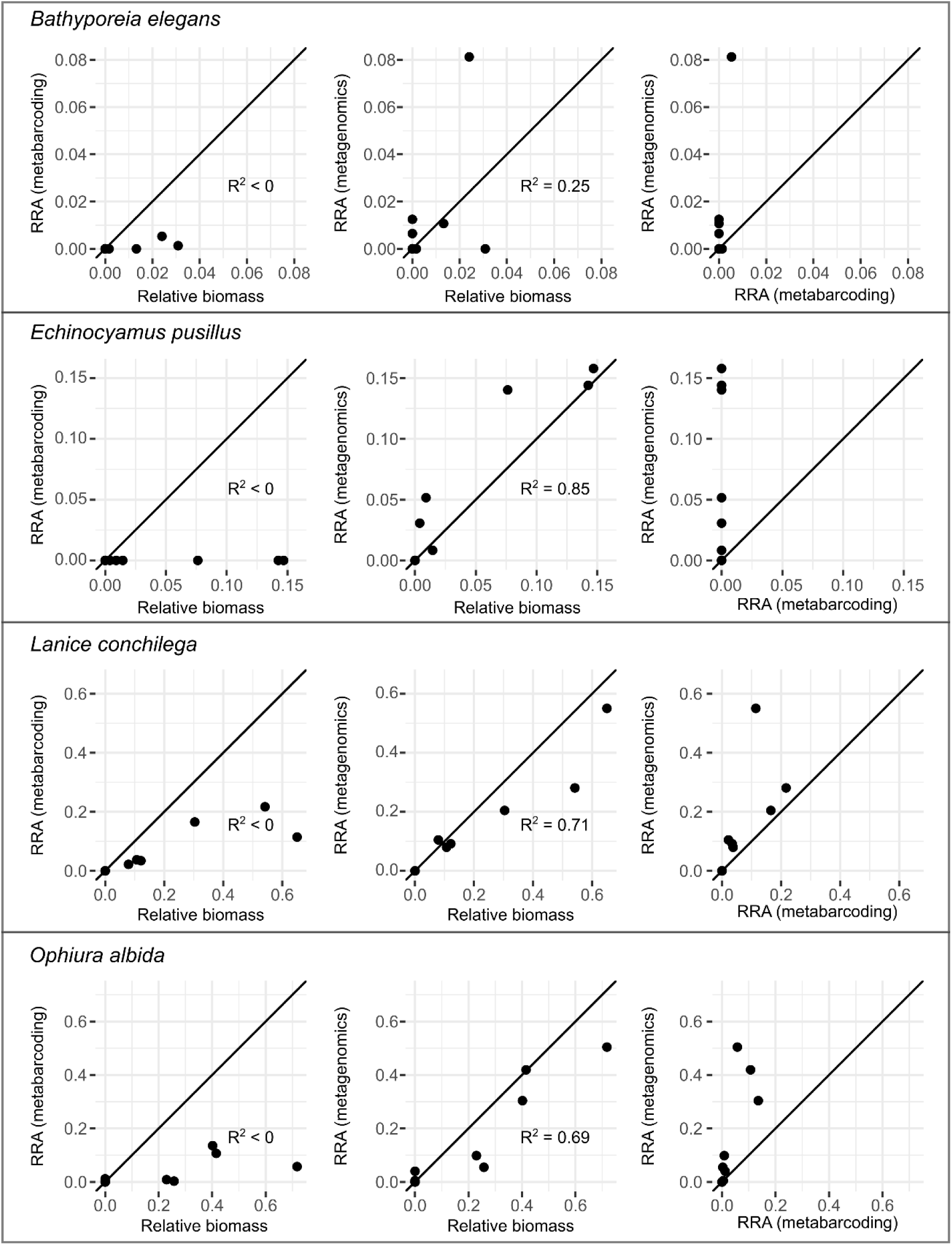
Species with indications that metagenomics alleviated a negative PCR bias. Left plot: Relation between relative read abundance (RRA) obtained with metabarcoding and relative biomass. Center plot: relation between RRA obtained with metagenomics and relative biomass. For both the left and center plots: points below the diagonal line indicate an under-estimation of relative biomass when using a DNA-based methods, points above the diagonal line indicate an over-estimation. Right plot: relation between RRA’s obtained with metagenomics and metabarcoding, points above the diagonal indicate that RRA obtained with metagenomics is higher than RRA obtained with metabarcoding. R^2^ values indicate the goodness-of-fit to the diagonal line.

**FIGURE 4.**
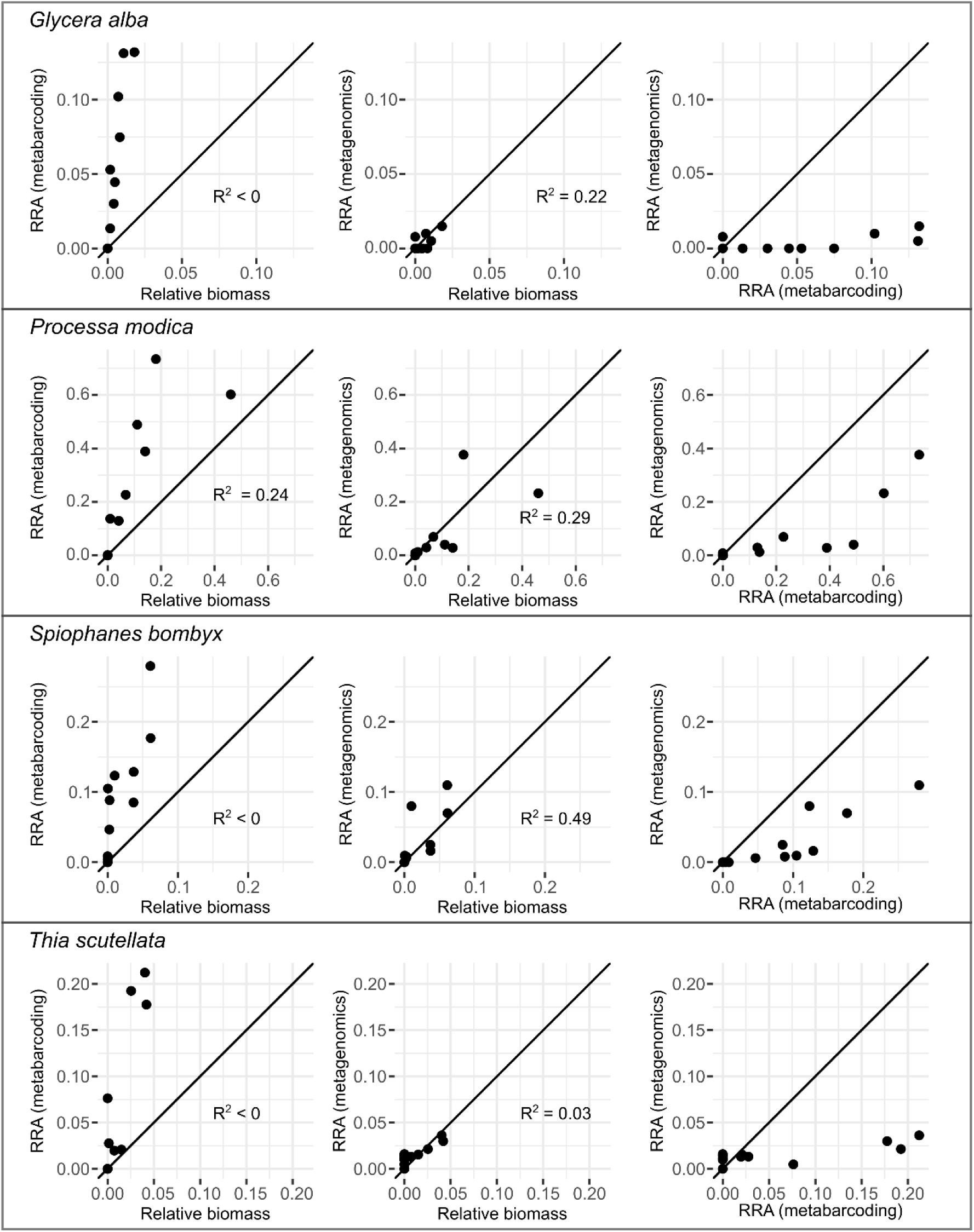
Species with indications that metagenomics alleviated a positive PCR bias. Left plot: Relation between relative read abundance (RRA) obtained with metabarcoding and relative biomass. Center plot: relation between RRA obtained with metagenomics and relative biomass. For both the left and center plots: points below the diagonal line indicate an under-estimation of relative biomass when using a DNA-based methods, points above the diagonal line indicate an over-estimation. Right plot: relation between RRA’s obtained with metagenomics and metabarcoding, points below the diagonal indicate that RRA obtained with metagenomics is lower than RRA obtained with metabarcoding. R^2^ values indicate the goodness-of-fit to the diagonal line.

### Estimations of required sequencing depth

#### Reference database

Figure 5 shows a clear and consistent relationship between the sequencing depth that was used to construct the reference database and the number of reads from metagenome samples that were classified. The reference database with lowest sequencing depth of 1 million reads already resulted in the classification of 42.91 ±0.16% and 45.92 ±0.04% of reads for *C. harengus* and *P. platessa*, respectively. The number of classified reads approached an asymptote for both species at around 6X genome coverage. Interestingly, the number of reads classified when using databases produced with >6X unassembled sequencing data slightly exceeded the number of reads classified using the benchmark database produced with an assembled reference genome (maximum of 102% for both species). The relative positions of the curves for both species indicate an effect of genome size.

**FIGURE 5.**
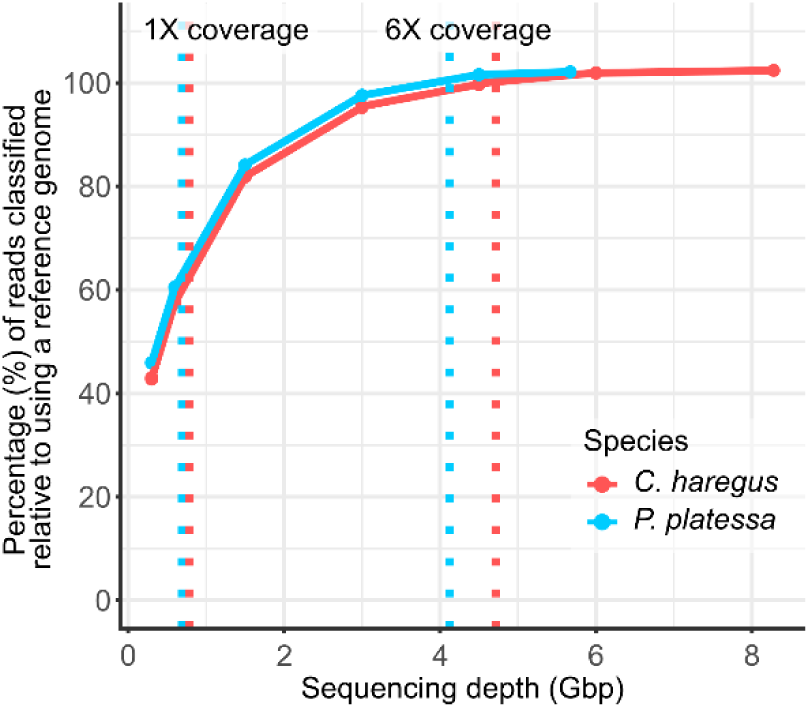
Impact of sequencing depth for reference database construction. The percentage of reads classified when using a database constructed with a given sequencing depth relative to using a benchmark database for two different fish species. The dotted lines indicate a sequencing depth of 1X and 6X coverage for each species.

#### Metagenomic sequencing

For each sample, the species detected at lower sequencing depths were already very similar to those obtained by the highest sequencing depth (Table 3; Figure 6A). Both missing species and additional species caused differences in diversity between subsets and full sequencing depth. The detection of additional species at lower sequencing depths was caused by a slight increase in their relative abundance, surpassing the 0.2% threshold. Additional species had a greater impact on the Jaccard similarity than missing species, with 80 instances of an additional species and 21 instances of a missing species over all subsets and replicates (supplementary Figure S3). The Bray-Curtis similarity between subsets of 10^3^ reads and full sequencing depth was already very high (for all replicate subsets Bray-Curtis similarity > 0.89; Table 3; Figure 6B), and community composition was practically identical for all samples when comparing subsets of 10^5^ reads to the full sequencing depth.

**FIGURE 6.**
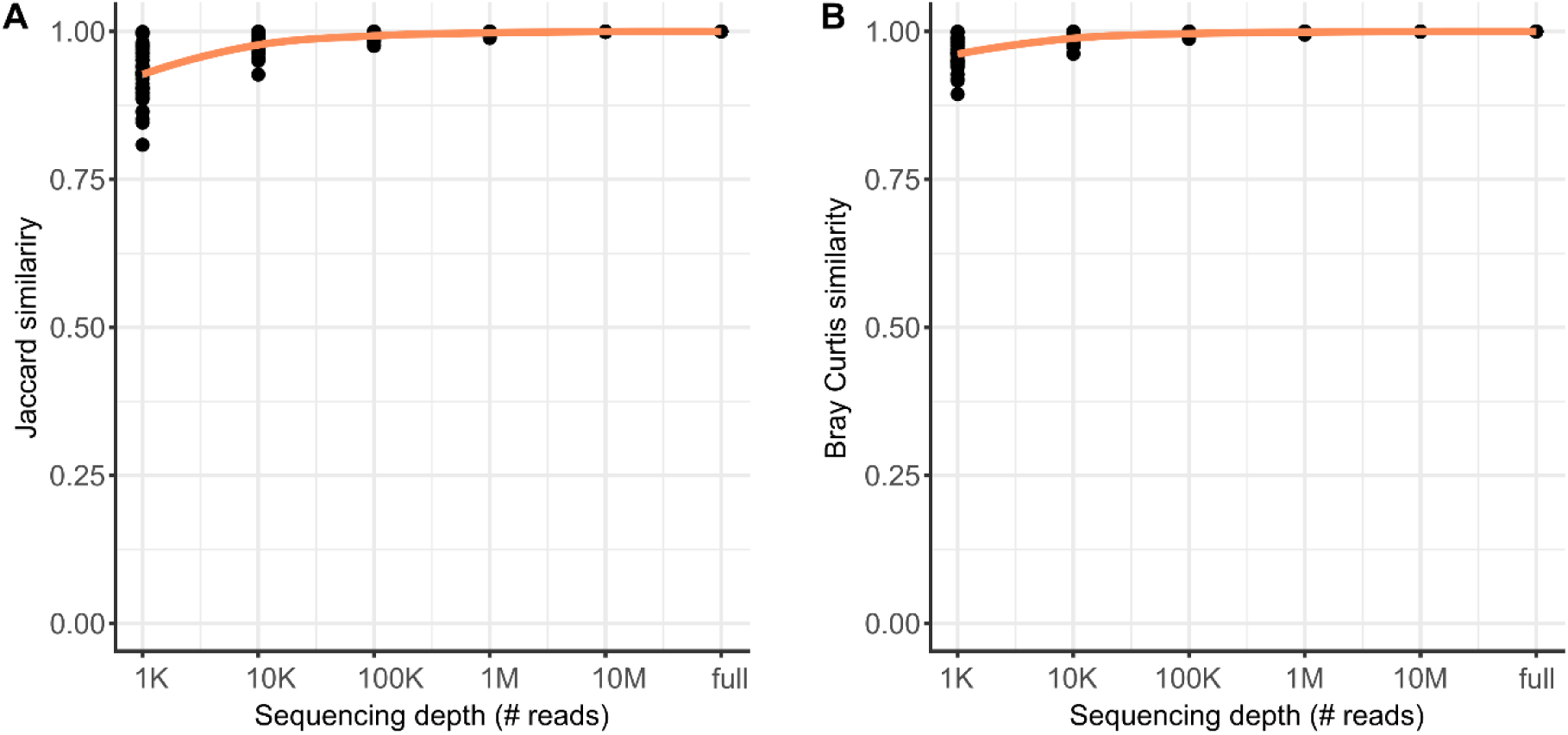
Estimation of required sequencing depth for metagenomic sequencing. Jaccard (A) and Bray-Curtis (B) similarity between community composition obtained with the full set of reads and subsets of reads with a range of sequencing depths (103 to 107) for each sample. Values for the ‘full’ dataset are included as visual reference, but are per definition 1.00 as the Jaccard and Bray-Curtis similarities were calculated relative to this full dataset.

**TABLE 3.** Jaccard similarity and Bray-Curtis similarity of community composition obtained with subsets of reads compared to the full sequencing depth of metagenomic samples.

| Sequencing depth<br>(# reads) | Jaccard similarity<br>(mean ± SD) | Bray-Curtis similarity<br>(mean ± SD) |
| --- | --- | --- |
| $10^3$ | $0.93 \pm 0.04$ | $0.96 \pm 0.02$ |
| $10^4$ | $0.98 \pm 0.02$ | $0.99 \pm 0.01$ |
| $10^5$ | $0.99 \pm 0.01$ | $1.00 \pm 0.00$ |
| $10^6$ | $1.00 \pm 0.00$ | $1.00 \pm 0.00$ |
| $10^7$ | $1.00 \pm 0.00$ | $1.00 \pm 0.00$ |

## DISCUSSION

The unavailability of reference genomes for most species in any ecosystem is one of the primary bottlenecks for the implementation of shotgun metagenomics for biodiversity assessments (Bell et al., 2021; Cowart et al., 2018; Garrido-Sanz et al., 2020; Schmidt et al., 2022; Serite et al., 2023b). In this study we developed a strategy using unassembled low-coverage whole genome sequencing data to build a reference database for metagenomic classification. Our results show that this is a cost-effective strategy to study community composition of bulk samples, with better estimates of relative abundances than the PCR-based metabarcoding method.

### Feasible whole-genome reference database construction for taxonomic assignment of metagenomic reads from diverse communities

In recent years, the generation of low-coverage whole genome sequencing data for biodiversity studies (i.e. genome skimming) has become commonplace for assembling high-copy genomic regions (mostly mtDNA and rDNA) with the goal of obtaining marker gene sequences or to perform phylogenetic analyses on a large number of species (Bista et al., 2018; Coissac et al., 2016; Hoban et al., 2022; Quattrini et al., 2024; Zhang et al., 2023). We show that sequencing data with a similar sequencing depth as genome skimming can also be used directly as reference data for shotgun metagenomic analysis, bypassing the need for assembly. In contrast to genome skimming, which often incorporates only around 1% of the generated sequencing data (Bohmann et al., 2020), our strategy incorporates virtually all the sequencing data that is generated for a species to construct a reference database. Yet, only a fraction of the required sequencing depth and limited bioinformatic processing are needed compared to e.g. the Earth BioGenome Project recommendations for generating reference genomes (https://www.earthbiogenome.org/). In addition, our strategy also incorporates sequencing data from regions which are difficult to assemble and are often missing from reference genomes such as repetitive rDNA arrays (Nurk et al., 2022). This might explain why slightly more metagenomic reads of *C. harengus* and *P. platessa* were classified using a database constructed from unassembled reads compared to using a database constructed with their reference genomes. A conceptually similar assembly-free approach has previously been developed for calculating the genomic distance between pairs of species, allowing identification of single specimens (Bohmann et al., 2020; Coissac et al., 2016; Sarmashghi et al., 2019). Our strategy builds further upon this assembly-free approach and extends it to relative abundance estimations in multi-species samples.

### Metagenomics improves biomass estimates of bulk macrobenthos samples

We hypothesized that shotgun metagenomics could provide a better characterization of macrobenthos communities than metabarcoding by removing the PCR-bias that we observed in earlier metabarcoding analyses (Derycke et al., 2021; Van den Bulcke et al., 2021, 2023, 2024). Our results show that metagenomics was superior to metabarcoding for approximating relative biomass based community composition. These improved relative abundance estimates could be attributed to two main causes: 1) the detection of high biomass species not detected by metabarcoding, and 2) an improved estimate of relative biomass for most species when compared to metabarcoding.

Detection of high biomass species not detected by metabarcoding was most apparent for the echinoderm *Echinocyamus pusillus*, for which the biomass could make up almost 15% of the community. This species remained undetected by metabarcoding, while metagenomics provided good estimates of its biomass. This seemed to be caused by the species-specific primer melting temperature, as none of the species with a minimal T_m_ below 50°C (including *E. pusillus*) was detected by metabarcoding. Bista et al. (2018) also showed that primer incompatibility was an important cause for bias in their metabarcoding data, with some high-biomass species remaining undetected. Their improved estimates of macroinvertebrate biomass when using mitochondrial metagenomics instead of metabarcoding could thus in part be attributed to additional detection of several high biomass species. Serite et al. (2023) also noted this cause for inducing a bias in dietary analysis of pipefishes when using metabarcoding, where key copepod species detected by metagenomics were not detected by metabarcoding.

Metagenomics also generally provided a better predictive value for relative biomass (higher R^2^) than metabarcoding for species that were detected by both methods, resulting in a better approximation of community composition. Especially the improved estimates for species that often obtain a high relative biomass, such as *Lanice conchilega*, *Ophiura albida* and *Nephtys cirrosa*, could have a large effect on Bray-Curtis similarity to biomass-based composition (Ricotta & Podani, 2017). Interestingly, Van den Bulcke et al. (2024) (analyzing the same samples) noted that *Thia scutellata* was one of the few species with a good correlation between metabarcoding read abundance and biomass. However, despite this good correlation, metabarcoding read abundance in our dataset did not provide a predictive value for relative biomass due to a consistent overestimation (Figure 3), resulting in a slope that strongly deviates from 1 when fitting a simple linear regression. In contrast, metagenomics read abundance did provide some predictive value for this species.

Although metagenomics generally performed better than metabarcoding for estimating relative biomass, metabarcoding showed better performance for two species: *E. cordatum* and *O. fusiformis*. For *E. cordatum,* we hypothesize that the biomass/DNA ratio is heavily skewed compared to other species because a large part of this animal’s biomass is made up of calcified material, introducing a strong bias independent of PCR. The reason for the consistent underestimation of *O. fusiformis* by metagenomics is not clear. In addition to the biomass/DNA ratio, other factors can also bias shotgun metagenomic estimates of relative biomass, such as variation in genome size (Chouvarine et al., 2016) or species-specific DNA extraction efficiency (McLaren et al., 2019). In future analyses, biases induced by genome size or the biomass/DNA ratio could potentially be accounted for if normalization factors are determined for each species.

Similar to Bista et al. (2018), we also observed false negatives of low-biomass species with both metagenomics and metabarcoding (respectively 15 and 28 cases in total). This was, for example, the case for *A. brachiata*, for which the relative biomass did not exceed 0.5% and remained undetected by both methods (although in sample 120, for which no biomass was recorded, this species was detected by morphology and metagenomics, but not by metabarcoding). In our metagenomic analysis, a lack of sequencing depth did not seem to be the reason for false negatives, as species detection remained relatively stable in subsamples where read numbers were decreased by an order of magnitude. The cause for false negatives in our analyses are the abundance thresholds we used for minimizing false positives. Drake et al. (2022) already indicated that setting abundance thresholds is a balancing act to minimize both false positives and false negatives. Indeed, removing our threshold led to a strong increase in false positives for both methods, while setting a threshold caused a small increase in false negatives, especially for metagenomics (Table 2). Other metagenomic studies similarly observed a high rate of false positives when not using some form of threshold (Garrido-Sanz et al., 2020; Schmidt et al., 2022). Although false positives frequently occur in metagenomic analyses, active algorithm development to better address this problem is ongoing (Breitwieser et al., 2018; Marini et al., 2024) and improvements might also lead to a better avoidance of false negatives when both sensitivity and specificity increase.

### Future perspectives

Metabarcoding can rely on a much larger reference database with over a million species represented in BOLD (Ratnasingham et al., 2024), while only 18 920 eukaryotic reference genomes are available on NCBI (as of December 2024). Undoubtedly, generating a substantial genomic database for any ecosystem will involve a considerable investment. Nonetheless, projects with this goal in mind are starting to emerge (e.g. the MetaInvert database for soil invertebrates; Collins et al., 2023). With the assembly-free approach we present here, genomic reference data for metagenomic classification can be generated for less than €100 per species and with limited bioinformatic processing. Therefore, the inclusion of several dozen of species from an ecosystem should be feasible within the time and budget of most scientific projects. In addition, species for which whole genome sequencing data is already publicly available (ranging from raw reads for genome skimming to assembled reference genomes) can readily be incorporated in custom databases.

We further show that with a sequencing effort similar to metabarcoding (e.g. Buchner et al., 2024) we obtained better estimates of community composition, making this a promising approach for large-scale biodiversity monitoring. However, given that this was only a limited study exploring the application of our approach, several aspects still need testing such as the ability to distinguish between closely related species, the impact and mitigation of bias introducing factors, and the performance on samples from a variety of ecosystems encompassing a wide range of diversity patterns. Nonetheless, the strategy presented here should be relatively easy to adopt for laboratories familiar with metabarcoding and can be used as an accessible alternative.

## ACKNOWLEDGEMENTS

We would like to thank Hans Hillewaert and Sara Maes for respectively providing images and DNA extracts of vouchered specimens. We would like to thank the crew of RV Belgica for assistance during sampling. This research was financed through the DNASense project, funded by the Biodiversa+ call with Belspo as national funder (Belspo grant number RT/24/DNASense_ILVO); FPS Economy through revenues of marine aggregate extraction in Belgian waters; and the Brexit Adjustment Reserve fund of the European Union (projectnumber: BAR0159).

## DATA AVAILABILITY

All sequencing data generated in this study is deposited on ENA project PRJEB83993.

