## Supplemental information for "An accessible metagenomic strategy allows for better characterization of invertebrate bulk samples"

### Supplementary materials

#### Vouchering images for species not available on BOLD

For other species, vouchering information can be found at our institutional portal on BOLD: https://portal.boldsystems.org/institution/Flanders%20Research%20Institute%20for%20Agriculture%2C%20Fisheries%20and%20Food

**_Nephtys caeca_ (ILVO002)**

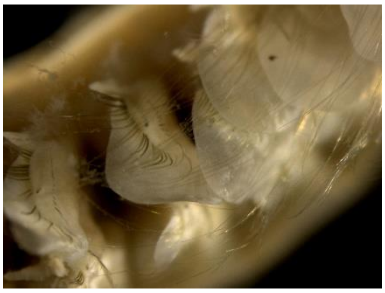

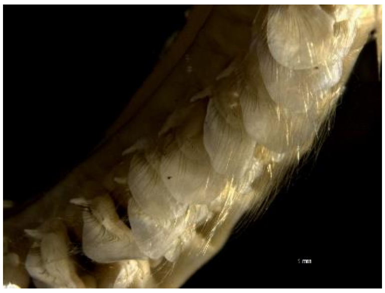

**_Nephtys longosetosa_ (ILVO003)**

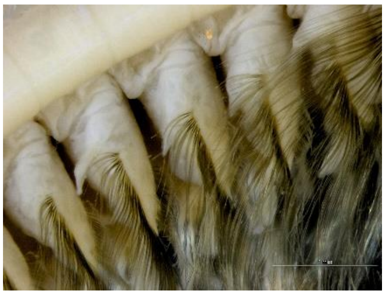

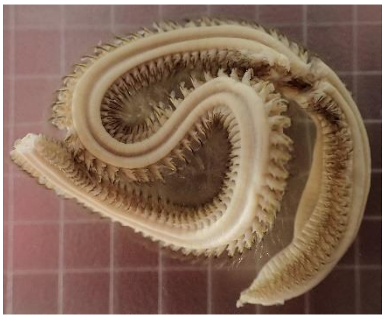

**_Glycera alba_ (ILVO018)**

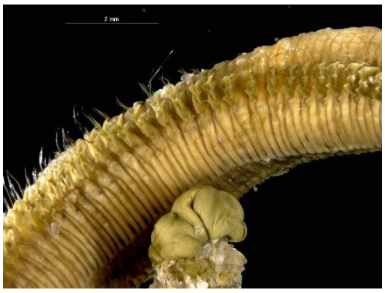

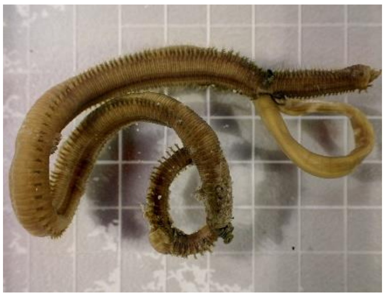

*Lanice conchilega* (ILVO475)

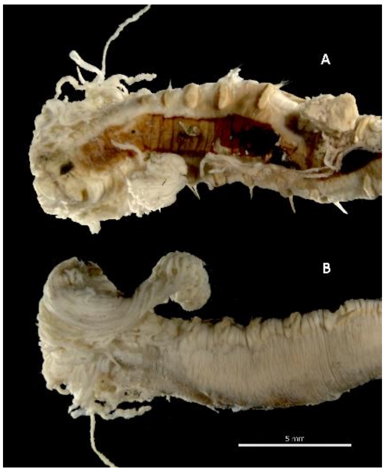


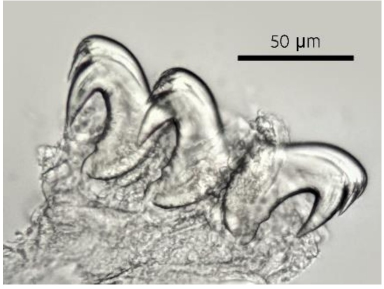


*Notomastus latericeus* (ILVO015)

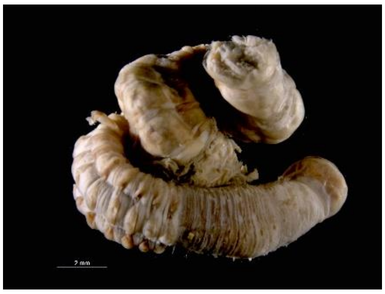

*Abra alba* (ILVO008)

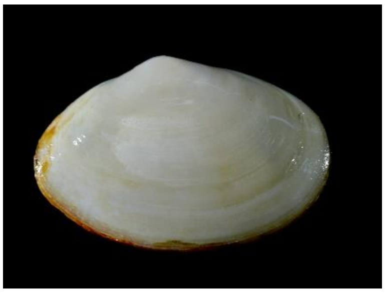

*Thia scutellata* (ILVO028)

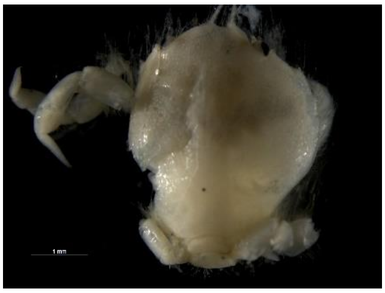

*Acrocnida brachiata* (ILVO223)

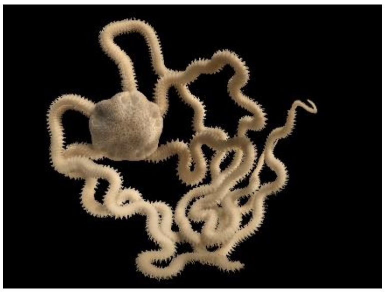

*Echinocyamus pusillus* (ILVO335)

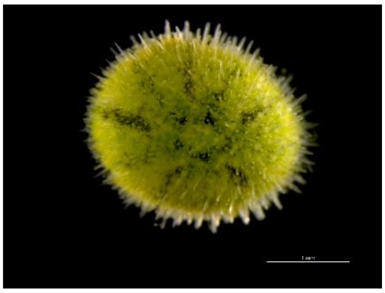

*Echinocardium cordatum* (ILVO012)

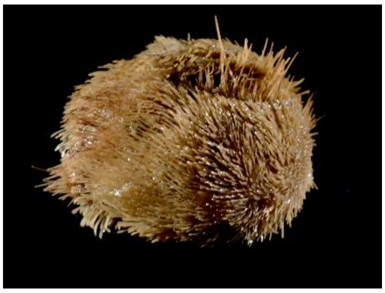

| Sample | BioSample accession | Sequencing depth (Million paired-end reads) |
|---|---|---|
| 120 | SAMN17039393 | 14.4 |
| ZVL | SAMN17039316 | 20.1 |
| TB11_19 | SAMN43265897 | 25.3 |
| TB12_19 | SAMN43265898 | 29.3 |
| TB12_21 | SAMN43265947 | 16.3 |
| TB16_21 | SAMN43265948 | 24.1 |
| TB47_21 | SAMN43265956 | 19.8 |
| TB48_19 | SAMN43265908 | 25.0 |
| TB48_21 | SAMN43265957 | 25.0 |
| TB54_19 | SAMN43265913 | 36.5 |
| TB54_21 | SAMN43265961 | 20.9 |
| TB56_19 | SAMN43265915 | 19.9 |
| TB59_19 | SAMN43265918 | 19.5 |
| TB66_21 | SAMN43265968 | 22.8 |

**Supplementary table S1:** List of metagenomic samples sequenced in this study. The BioSample links to both the metabarcoding and metagenomic read files used for our analysis. The BioSample also lists all sample metadata. The sequencing depth (in million paired-end reads) is provided for the shotgun metagenomic run. Details on sequencing data for the metabarcoding run can be found in Van den Bulcke et al. (2021) and Van den Bulcke et al. (2024).

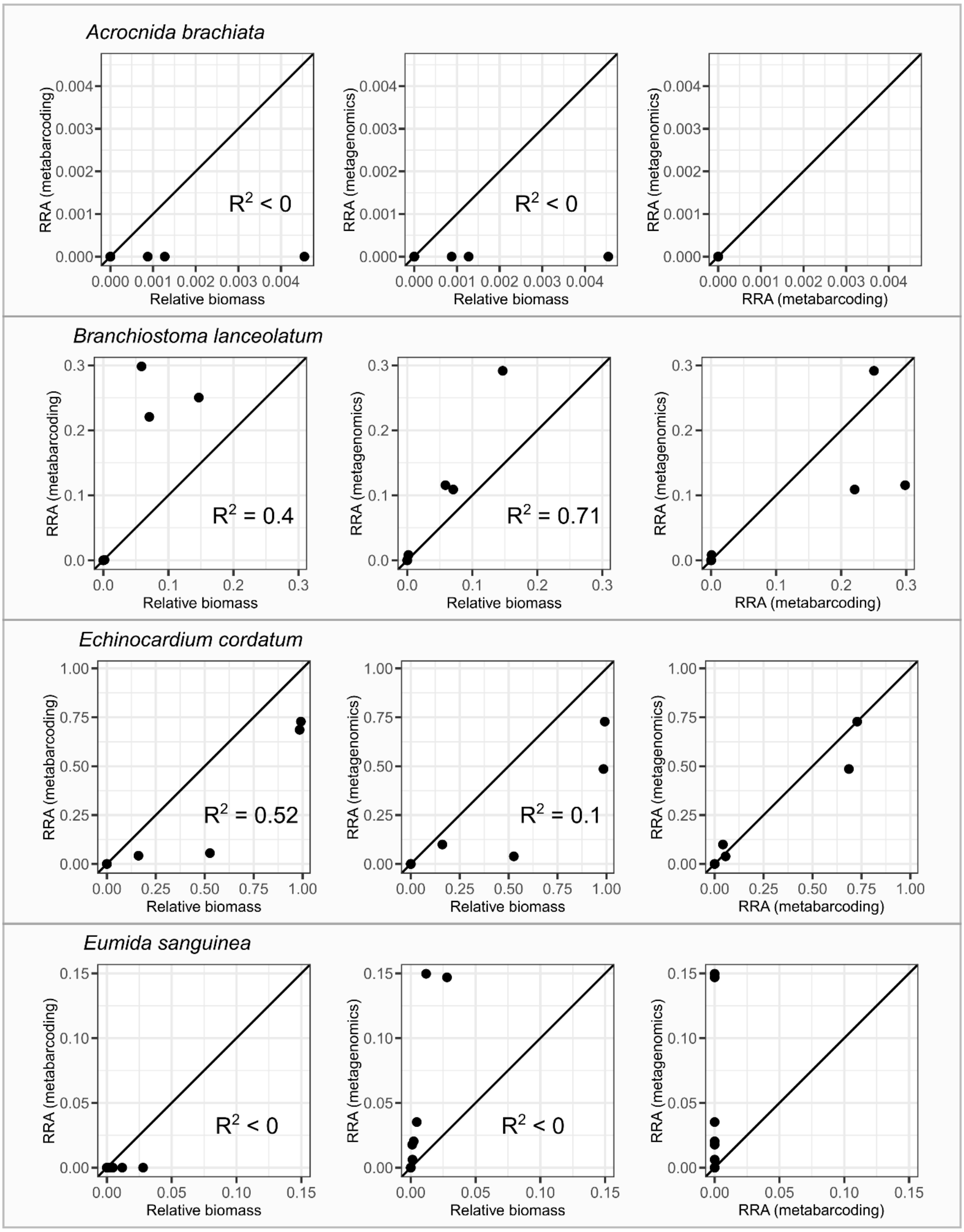


**Supplementary Figure S1**: Left plot: Relation between relative read abundance (RRA) obtained with metabarcoding and relative biomass. Center plot: relation between RRA obtained with metagenomics and relative biomass. Right plot: relation between RRA's obtained with metagenomics and metabarcoding. $R^2$ values indicate the goodness-of-fit to the diagonal line.

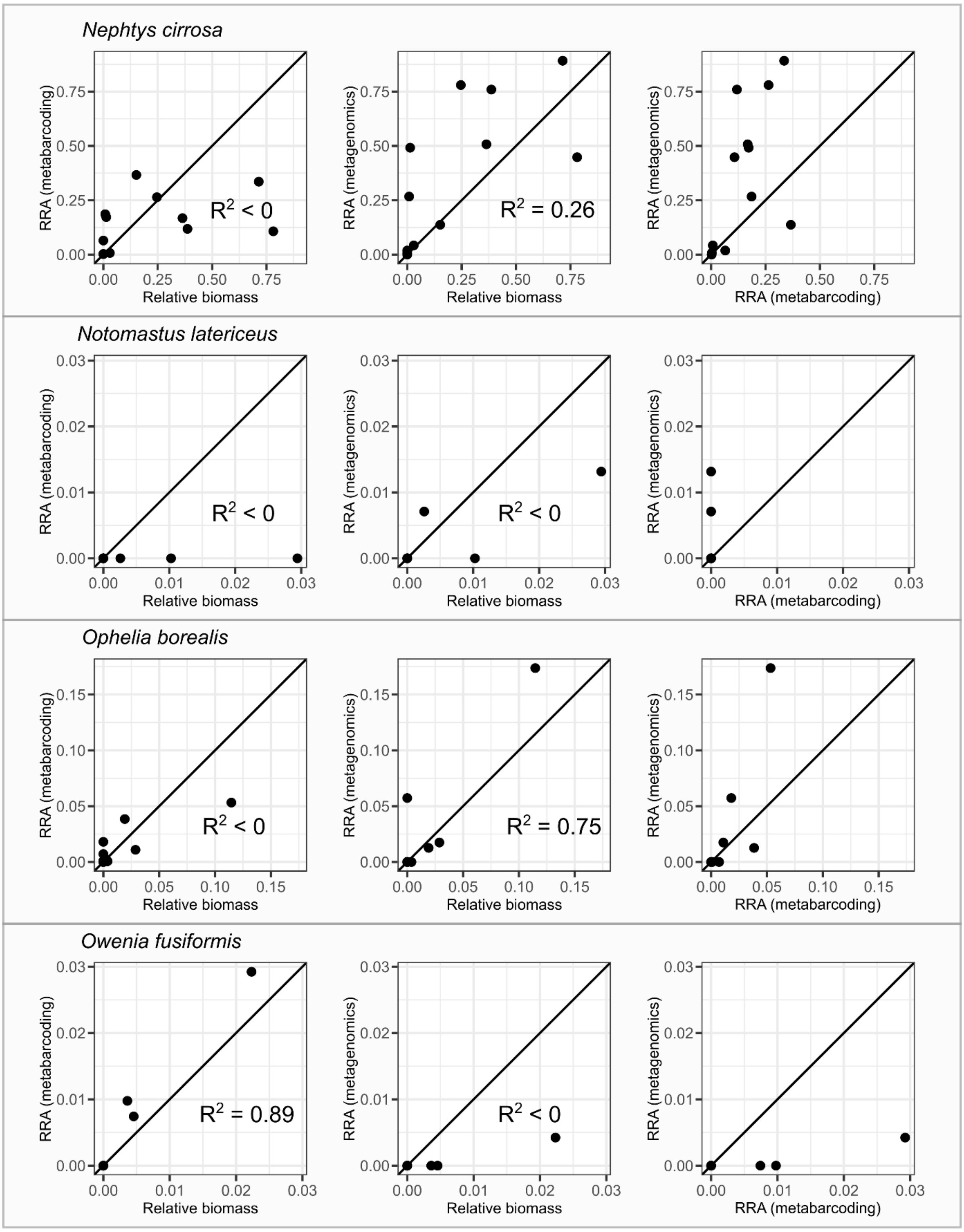


**Supplementary Figure S2**: Left plot: Relation between relative read abundance (RRA) obtained with metabarcoding and relative biomass. Center plot: relation between RRA obtained with metagenomics and relative biomass. Right plot: relation between RRA's obtained with metagenomics and metabarcoding. $R^2$ values indicate the goodness-of-fit to the diagonal line.

**Supplementary Table S2:** Melting temperatures for metabarcoding primers (5'-3') mICOIintF (GGWACWGGWTGAACWGTWTAYCCYCC) and jgHCO2198 (TANACYTCNGGRTGNCCRAARAAYCA) calculated for each species using their assembled mitochondrial genomes as template. Melting temperatures were obtained using SnapGene Viewer 7.1. 'NA' indicates that no primer binding site was found for a particular primer on the mitochondrial assemblies. Species for which no metabarcoding reads were obtained are indicated in red.

| Species | $T_m$ (mICOIintF) | $T_m$ (jgHCO2198) | Detection by metabarcoding |
|---|---|---|---|
| ***Eumida sanguinea*** | 41°C | 62°C | No |
| ***Nephtys cirrosa*** | 57°C | 59°C | Yes |
| ***Nephtys hombergii*** | 54°C | 60°C | Yes |
| ***Glycera alba*** | NA | 56°C | Yes |
| ***Magelona johnstoni*** | 60°C | 59°C | Yes |
| ***Scolelepis bonnieri*** | 60°C | 61°C | Yes |
| ***Spiophanes bombyx*** | 61°C | 64°C | Yes |
| ***Owenia fusiformis*** | 59°C | 58°C | Yes |
| ***Lanice conchilega*** | 53°C | 55°C | Yes |
| ***Notomastus latericeus*** | 46°C | 58°C | No |
| ***Ophelia borealis*** | NA | 63°C | Yes |
| ***Abra alba*** | 40°C | 62°C | No |
| ***Macoma balthica*** | 58°C | 57°C | Yes |
| ***Fabulina fabula*** | NA | 57°C | No |
| ***Bathyporeia elegans*** | 53°C | 55°C | Yes |
| ***Processa modica*** | 61°C | 62°C | Yes |
| ***Liocarcinus depurator*** | 58°C | 62°C | Yes |
| ***Thia scutellata*** | 57°C | 59°C | Yes |
| ***Ophiura ophiura*** | 61°C | 56°C | Yes |
| ***Ophiura albida*** | 57°C | 57°C | Yes |
| ***Acrocnida brachiata*** | 63°C | 60°C | No |
| ***Echinocyamus pusillus*** | 42°C | 58°C | No |
| ***Echinocardium cordatum*** | 50°C | 61°C | Yes |
| ***Branchiostoma lanceolatum*** | 51°C | 56°C | Yes |

**Supplementary Figure S3:** Species detection using morphology and metagenomics across all stations. Each subplot corresponds to a sampling station, with species listed on the y-axis and replicates for each subsample size ($10^3$ to $10^7$) and the full set of reads shown on the x-axis. Dots are colored according to the species detection in each replicate with metagenomics (red), morphology (blue), both methods (green) and none (grey).

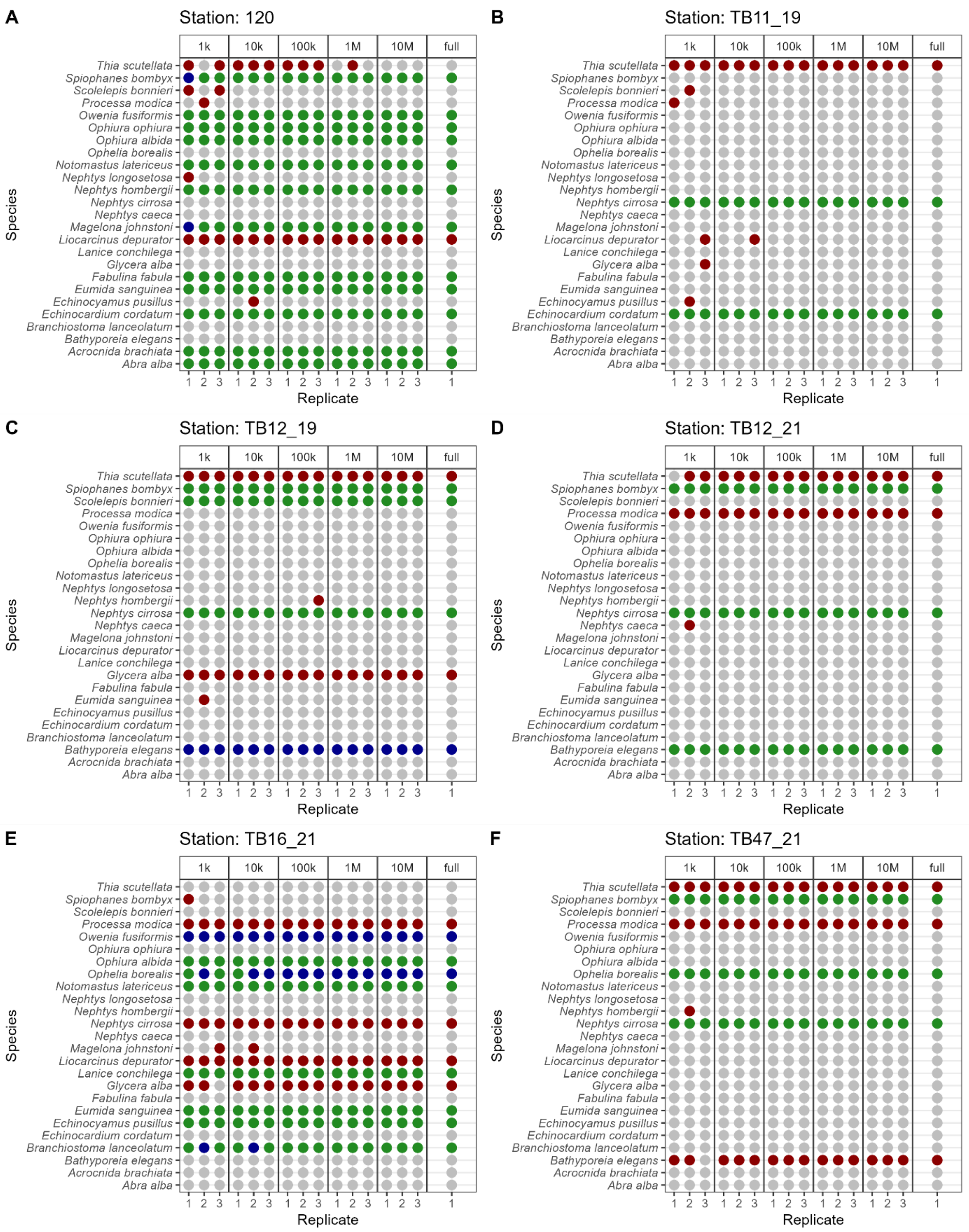

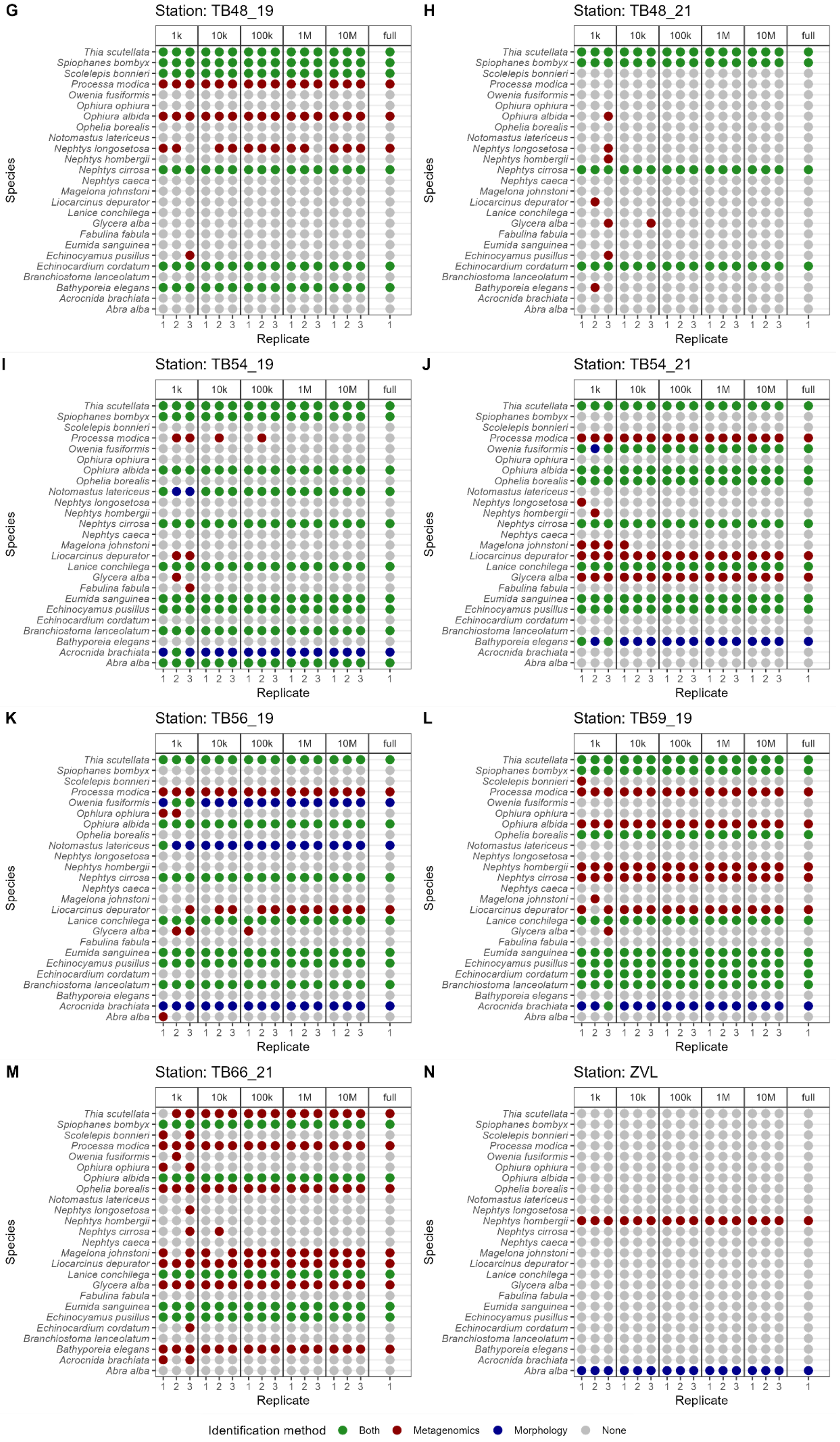
G
Station: TB48_19
H
Station: TB48_21
I
Station: TB54_19
J
Station: TB54_21
K
Station: TB56_19
L
Station: TB59_19
M
Station: TB66_21
N
Station: ZVL
1k
10k
100k
1M
10M
full
Thia scutellata
Spiophanes bombyx
Scolelepis bonnieri
Processa modica
Owenia fusiformis
Ophiura ophiura
Ophiura albida
Ophelia borealis
Notomastus latericeus
Nephtys longosetosa
Nephtys hombergii
Nephtys cirrosa
Nephtys caeca
Magelona johnstoni
Liocarcinus depurator
Lanice conchilega
Glycera alba
Fabulina fabula
Eumida sanguinea
Echinocyamus pusillus
Echinocardium cordatum
Branchiostoma lanceolatum
Bathyporeia elegans
Acrocnida brachiata
Abra alba
Species
1 2 3
Replicate
Identification method
Both
Metagenomics
Morphology
None

### Code for bioinformatic and statistical analysis

#### Bioinformatic processing of reference sequence data

The code below shows the methodology used genomic reference database construction, resulting in a custom kraken2 k-mer index database (top panel of Figure 1 “Genomic reference database construction”)

#### Pre-processing

Pre-processing includes read QC through visual inspection, trimming low quality bases, merging overlapping paired-end reads, and combining all sequencing data in a single fastq file. The workflow is shown for a single species (with files species_1.fq and species_2.fq being the forward and reverse reads produced by an 2x150bp Illumina run) and should be done for each species separately.

##### *readQC*

Run fastQC

```
fastqc -threads 16 species_R1.fastq species_R2.fastq -o fastqc_output_dir
```

Navigate to the directory containing the fastQC output

```
cd fastqc_output_dir
```

Combine fastQC results into a single html file

```
multiqc .
```

##### *Trimmomatic*

Remove low quality bases and ILLUMINA nextera adapters from reads. This produces four files:

- species_paired_R1.fastq (trimmed, paired forward reads)
- species_unpaired_R1.fastq (trimmed, unpaired forward reads)
- species_paired_R2.fastq (trimmed, paired reverse reads)
- species.unpaired_R2.fastq (trimmed, unpaired reverse reads)

```
trimmomatic PE \
-threads 8 \
species_R1.fastq species_R2.fastq \
species_paired_R1.fastq species_unpaired_R1.fastq species_paired_R2.fastq
species_unpaired_R2.fastq \
ILLUMINACLIP:adapters/NexteraPE-PE.fa:2:30:10 \
HEADCROP:3 \
TRAILING:3 \
SLIDINGWINDOW:4:30 \
```

##### *FastqJoin*

The paired reads (species_paired_R1.fastq and species_paired_R2.fastq files) are merged using FastqJoin, which produces three files:

- species_paired_merged.fastq (merged read pairs)
- species_paired_unmerged_R1.fastq (forward read of non-overlapping reads pairs)
- species_paired_unmerged_R2.fastq (reverse read of non-overlapping read pairs)

```
fastq-join species_paired_R1.fastq species_paired_R2.fastq \
-o species_paired_unmerged_R1.fastq -o species_paired_unmerged_R2 -o
species_paired_merged.fastq
```

*Combine in a single fastq file*

All the reads (unpaired R1 and R2, paired unmerged R1 and R2, and merged) are now combined into a single fastq file (species_all_reads_concat.fastq).

```
cat species_paired_merged.fastq species_paired_unmerged_R1.fastq
species_paired_unmerged_R2.fastq species_unpaired_R1.fastq
species_unpaired_R2.fastq > species_all_reads_concat.fastq
```

#### Contamination filtering

This step uses the standard kraken2 index to classify potential bacterial, viral and human reads. The latest version of the standard kraken2 database can be downloaded from https://benlangmead.github.io/aws-indexes/k2.

The first step classifies all reads in the species_all_reads_concat.fasta file using the kraken2 standard database. This produces the ClassifiedContamination.kraken file.

```
kraken2 --use-names --threads 8 --db standard --report
ReportContamination.txt species_all_reads_concat.fastq >
ClassifiedContamination.kraken
```

The next step uses KrakenTools/extract_kraken_reads.py at master · jenniferlu717/KrakenTools · GitHub to extract all the unclassified reads from the species_all_reads_concat.fastq file, which produces the species_filtered.fasta file.

```
python3 extract_kraken_reads.py -k ClassifiedContamination.kraken -s
species_all_reads_concat.fastq -t 0 -o species_filtered.fasta
```

#### Reference database construction

Instead of assembling the reads, we will use the unassembled reads directly to construct a reference database. The first step is to add the taxid of your species to each sequence header in the species_filtered.fasta file. We wrote a custom python script (Add_Taxid.py) to accomplish this:

```
import argparse, re
parser = argparse.ArgumentParser(description='Add the Taxonomy to the
sequence headers')
parser.add_argument('inputFile', type=str,
                    help='Provide the input file to add the taxid ID to the
sequence headers')
parser.add_argument('outputFile', type=str,
                    help='Provide the (path/)name for the output file')
parser.add_argument('-t', '--taxid', type=str, required = True,
                    help ='The taxonomy ID for the reads provided.')
args = parser.parse_args()
with open(args.outputFile, "w") as file_to_write:
    with open(args.inputFile, "r") as file_to_read:
```

```
        file_lines = file_to_read.readlines()

        for line in file_lines:
            if re.search(">", line):
                stripped_line = line.strip()
                seq_taxid = ">kraken:taxid|" + args.taxid + "|" +
stripped_line[1:] + "\n"
                file_to_write.writelines(seq_taxid)
            else:
                file_to_write.writelines(line)
    file_to_read.close()
file_to_write.close()
```

To run this script from the command line (assuming the species taxid = 123456):

```
python3 Add_Taxid.py -t 123456 species_filtered.fasta 123456_species.fasta
```

All of the processing above should be done for each species separately, resulting in a single fasta file with name structure taxid_speciesName.fasta. The fasta files of all species included in the reference database should then be placed in a single directory (in this example named "species_fasta_lib").

The next step add all the fasta files to the kraken2 library to start constructing the k-mer index database (here named 'customDB')

```
for file in species_fasta_lib/*.fasta; do
kraken2-build --threads 16 --add-to-library $file --db customDB
done
```

Then download the NCBI taxonomy:

```
kraken2-build --threads 16 --use-ftp --download-taxonomy --db customDB
```

Finally, build the database

```
kraken2-build --threads 8 --build --db customDB
```

#### Bioinformatic processing of metagenomic sequencing data

Metagenomic classification to obtain a species abundance table from metagenomic sequencing data (forward and reverse reads from an illumina run, here named community_R1.fastq and community_R2.fastq) involves three steps: readQC/trimming, metagenomic classification, and conversion of the kraken report to biom format (which can be imported in e.g. for statistical analysis

##### ReadQC/Trimming

*readQC*

Run fastQC

```
fastqc -threads 16 community_R1.fastq community_R2.fastq -o
fastqc_output_dir
```

Navigate to the directory containing the fastQC output

```
cd fastqc_output_dir
```

Combine fastQC results into a single html file

```
multiqc .
```

##### *Trimmomatic*

Remove low quality bases and ILLUMINA nextera adapters from reads. This produces four files:

- community_paired_R1.fastq (trimmed, paired forward reads)
- community_unpaired_R1.fastq (trimmed, unpaired forward reads)
- community_paired_R2.fastq (trimmed, paired reverse reads)
- community.unpaired_R2.fastq (trimmed, unpaired reverse reads)

```
trimmomatic PE \
-threads 8 \
community_R1.fastq community_R2.fastq \
community_paired_R1.fastq community_unpaired_R1.fastq
community_paired_R2.fastq community_unpaired_R2.fastq \
ILLUMINACLIP:adapters/NexteraPE-PE.fa:2:30:10 \
HEADCROP:3 \
TRAILING:3 \
SLIDINGWINDOW:4:15 \
```

*note : we have found that the trimming step is not strictly necessary when readQC shows good sequencing quality as shown in the plot below:

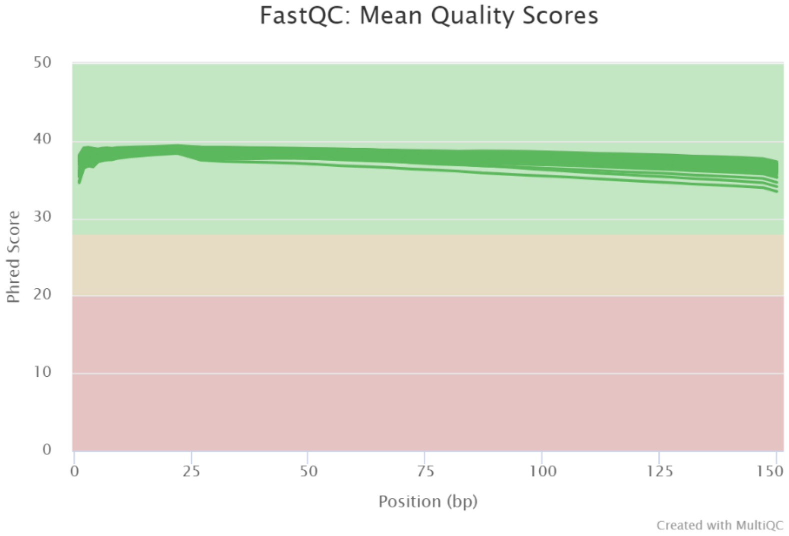


#### Metagenomic classification

We will use the "customDB" produced in the previous section to classify the metagenomic reads. Given that we use the --paired option, only reads from the community_paired_R1.fastq and community_paired_R2.fastq files are classified.

```
kraken2 --paired --output community.output.kraken --report
community.report.kraken --threads 16 --memory-mapping --minimum-hit-groups
3 --db customDB community_paired_R1.fastq community_paired_R2.fastq
```

#### Convert to biom format and import in phyloseq

This is optional, but for those who want to analyze their data in R using the phyloseq package, the following method converts the kraken2 reports to a biom format, which can be imported into R:

After classifying all your metagenomic samples, make sure the reports are in the same directory.

Install kraken-biom from https://github.com/smdabdoub/kraken-biom

Run from within the directory containing the report files, make sure to select the json format as phyloseq does not support hdf5. This will produce a file named metagenomic_classifications.biom

```
kraken-biom -o metagenomic_classifications.biom --fmt json
```

To import in R, load the phyloseq package, import the biom file and merge with the metadata to a phyloseq object

```
#Load the biom file with kraken classifications
biom <- import_biom(BIOMfilename = "metagenomic_classification.biom")

#Load the metadata and format for phyloseq
metadata <- read.csv("metadata.txt", sep = "\t")
rownames(metadata) <- metadata[,1]
sampledata <- sample_data(metadata)

#Remove the .., f__, s__ annotation from taxon names and change names of
taxonomic levels
biom@tax_table@.Data <- substring(biom@tax_table@.Data, 4)
colnames(biom@tax_table@.Data) <- c("Kingdom", "Phylum", "Class", "Order",
"Family", "Genus", "Species")

#Merge with sampledata
Metagenomic_data <- merge_phyloseq(biom, sampledata)
```
